# Comparative analysis of 15 chromosome-scale near T2T assemblies of Brazilian *Fusarium graminearum* isolates

**DOI:** 10.64898/2026.09.16.752100

**Authors:** Erika Kroll, Ana Machado Wood, Camilla P Nicolli, Arne De Klerk, Martin Urban, Emerson Del Ponte, Keywan Hassani-Pak, Kim E Hammond-Kosack

## Abstract

*Fusarium graminearum* is the causative agent of Fusarium head blight of wheat, yet genomic resources for Brazilian populations, where it is the dominant species, remain limited to a single reference genome. We report fifteen near telomere-to-telomere (T2T) assemblies of Brazilian *F. graminearum* isolates generated with Oxford Nanopore sequencing. Each genome was assembled into four nuclear chromosomes, with 118 of 120 telomeres resolved, BUSCO completeness of 99.1–99.3%, and 14,802–14,894 predicted genes per isolate. The genomes are structurally conserved, showing no chromosome-scale rearrangements and uniform transposable element content. Orthology assignment placed 90.4% of genes in the core genome. This suggests that the large phenotypic variance in aggressiveness observed between isolates likely resides in regulatory differences and/or within the small and largely uncharacterised accessory fraction. Ultimately, these assemblies provide a high-resolution resource for the global FHB community and a foundation for comparative analysis of a previously underrepresented population.

## Introduction

Fusarium head blight (FHB) is a major floral disease of wheat and other small grain cereals, including barley, maize, and oat, causing significant yield losses and reduced grain quality. FHB is caused by up to 17 Fusarium species, with *Fusarium graminearum sensu stricto* being the most predominant worldwide (Armer, Kroll, et al. 2024). *F. graminearum* produces the mycotoxin deoxynivalenol (DON), which is important to virulence and poses risks to human, animal, and ecosystem health. Reflecting its economic importance, *F. graminearum* was among the first filamentous fungal pathogens sequenced (Cuomo et al. 2007); the reference genome of the North American strain PH-1 was assembled into four chromosomes, later completed with centromeric and telomeric regions, and 14,164 gene models predicted (King et al. 2015; King et al. 2017).

While *F. graminearum* isolates from North America, Australia, Europe, Asia, and Brazil have been sequenced, earlier studies typically focused on single genomes or small isolate numbers (Cuomo et al. 2007; Gardiner et al. 2014; Walkowiak et al. 2016; Wang et al. 2017; Wood et al. 2020). Population genomic analysis of 60 North American isolates later revealed signatures of divergent evolution, including structure linked to the emergent NX-2 mycotoxin chemotype (Kelly and Ward 2018). More recently, analyses of 96 European strains revealed substantial geographic population structure (Kulik et al. 2023). Comparison of four nanopore-based chromosome-level assemblies of US isolates, has since shown that inversions, translocations and duplications concentrate near chromosome ends and in high-recombination regions (Dhakal et al. 2024).

Building upon foundational comparative genomics techniques, the field of pangenomics emerged as a framework to represent genomic variation at scale. A pangenome encompasses all genomes of a species, comprising a core genome shared by all individuals and an accessory genome of non-universal genes (Tettelin et al. 2005). Species level pangenomes are now available for several cereal pathogens, including *Zymoseptoria tritici* (Plissonneau et al. 2018; Chen et al. 2023), *Pyrenophora tritici-repentis* (Moolhuijzen et al. 2022), *Parastagonospora nodorum* (Richards et al. 2018), and *Pyrenophora teres* f. sp. *teres* (Wyatt et al. 2020). In *F. graminearum,* a pangenome built from primarily European isolates expanded the known gene repertoire by 32%, with only 53% of secreted protein clusters shared across isolates (Alouane et al. 2021). However, Brazilian *F. graminearum* isolates remain represented by a single genome assembly (Wood et al. 2020). Capturing the Brazilian pangenome is particularly important as *F. graminearum* is the dominant FHB-causing species in the country (de Chaves et al. 2022).

Here we close this gap by generating 15 chromosome-scale, near telomere-to-telomere (T2T) nanopore assemblies of *F. graminearum* isolates from southern Brazil. Long-read nanopore sequencing now enables near-complete, chromosome-scale assemblies with superior resolution of structural variation, repetitive regions, and accessory elements compared to short-read approaches, establishing T2T assemblies as the emerging gold standard (Li and Durbin 2024). By functionally annotating these genomes, and conducting a pangenomic analysis, we provide critical geographic and phenotypic data as a foundational resource for the global FHB community.

## Methods

### Origin, maintenance of Fusarium strains and DNA preparation

*F. graminearum* PH-1 and fifteen *F. graminearum* Brazilian *Coleção Micológica de Lavras* (CML) field isolates collected from wheat fields (2007-2011) are available from the Fungal Genetics Stock Center, Kansas City, MO, USA. *F. graminearum* isolates were grown in potato dextrose broth (PDB) or yeast extract peptone dextrose (YPD) liquid culture, with shaking at 100 rpm at 25°C for 3 days.

### Infection assay

Point-inoculations of wheat spikes cv. Bobwhite, were done at the first appearance of anther extrusion, using 5μl of 5 x 10^5^ conidia/ml spores. The *F. graminearum* conidial suspension was placed in the floral cavity between the palea and lemma of the outer two spikelets in the mid-region of the spike. The 13th and 14th spikelets (counting from the bottom of the spike) were point inoculated in each case. Six spikes were inoculated per isolate.

Inoculated plants were placed inside transparent boxes to retain high humidity for the first 48h (with the first 24h in the dark) and were then returned to 60% relative humidity for up to 20 days. Disease progress was recorded by counting the number of visibly diseased spikelets below the inoculation points on each wheat spike. Macroscopic disease symptoms were carefully monitored after fungal inoculation and scored at 15 days post infection (dpi).

### DNA extraction and whole genome sequencing

Genomic DNA was extracted from fungal mycelium using the Macherey-Nagel NucleoBond HMW DNA kit, with modifications optimised for *F. graminearum*. Isolates were cultured in 100 mL Yeast Peptone Sucrose (YPS) medium (0.6% peptone, 0.6% yeast extract, 2% sucrose, autoclaved separately) in siliconised 500 mL Erlenmeyer flasks, inoculated with 50 µL of a 5 × 10⁶ spores/mL suspension, and incubated at 28°C with shaking at 180 rpm for 2–3 days until the early exponential phase to minimise melanin accumulation. Mycelium was harvested by filtration or centrifugation, washed with sterile water, blotted dry (up to 2.5 g wet weight), and stored at −80°C prior to extraction. For each extraction, 0.8–1 g of mycelium was ground under liquid nitrogen with a mortar and pestle and processed following the manufacturer’s protocol, with RNA removed by digestion with 80 µL of 20 mg/mL RNase A (Invitrogen™ PureLink™) for 30 minutes at 37°C, rather than the 5-minute room-temperature step specified by the kit. DNA was resuspended in up to 250 µL EB buffer (Qiagen) depending on the pellet size.

### Nanopore sequencing and basecalling

High-molecular-weight genomic DNA from each of the 15 Brazilian *Fusarium graminearum* isolates was sequenced on an Oxford Nanopore PromethION using R10.4.1 flow cells (FLO-PRO114M) with libraries prepared using the Native Barcoding Kit 24 (SQK-NBD114-24). Reads were basecalled and demultiplexed within MinKNOW (v24.11.11; Bream v8.2.5, Configuration v6.2.12, MinKNOW Core v6.2.8) using the integrated Oxford Nanopore Dorado basecaller (v7.6.8); adapter and barcode sequences were removed during basecalling.

### De novo genome assembly

Basecalled reads were length-filtered with SeqKit (v2.2.0) (Shen et al. 2024) to retain reads ≥10 kb. Length-filtered reads were assembled using two complementary approaches: (i) directly with hifiasm (v0.19.7) in Nanopore mode (--ont --write-ec) (Cheng et al. 2026), and (ii) following HERRO error-correction in Dorado (v0.7.2; dorado correct, model herro-v1) (Stanojević et al. 2026), with the corrected reads then assembled using hifiasm (v0.19.7).

Contigs were assigned to chromosomes by aligning each assembly to the chromosome-level *F. graminearum* CML3066 v2 (Wood et al. 2020) reference with minimap2 (v2.26) (Li 2018) and visualising the alignments as dot plots (paf2dotplot); this identified which contigs corresponded to each of the four chromosomes, together with their relative order and orientation. Telomeric repeats were identified with tidk (v0.2.65) (Brown et al. 2025) (canonical TTAGGG array). In the absence of a full telomere-to-telomere chromosome contig, where possible the candidate contigs for that chromosome, selected from the dot plots, were self-aligned with minimap2 (-x asm5) to locate overlapping, telomere-bearing contig ends, which were then joined by extracting and concatenating the relevant coordinate intervals with SAMtools (v1.18) (Danecek et al. 2021).

### Mitochondrial genome assembly

Mitochondrial genomes were assembled with MitoHiFi (v. 3.2.2) (Uliano-Silva et al. 2023) from the length-filtered reads, using the *Fusarium graminearum* str. CML3066 mitochondrial genome (GenBank LT222057.1) as the seed reference. For each isolate the candidate best matching the reference in length and coverage was retained. Mitochondrial circularisation was assessed from the MitoHiFi output.

### Assembly quality assessment

Assembly completeness was evaluated with BUSCO (v. 5.7.1, lineage [hypocreales_odb10 / sordariomycetes_odb10]) (Manni et al. 2021), contiguity and size with QUAST (v 5.2.0) (Gurevich et al. 2013), and per-base consensus accuracy (QV) with yak (v.0.1-r69) (https://github.com/lh3/yak) using the corrected ONT reads. Genome coverage was calculated from the length-filtered reads against each final assembly with SeqKit v.2.2.0 (Shen et al. 2024). Assembly statistics for all 15 isolates are summarised in Table 1.

**Table 1.**
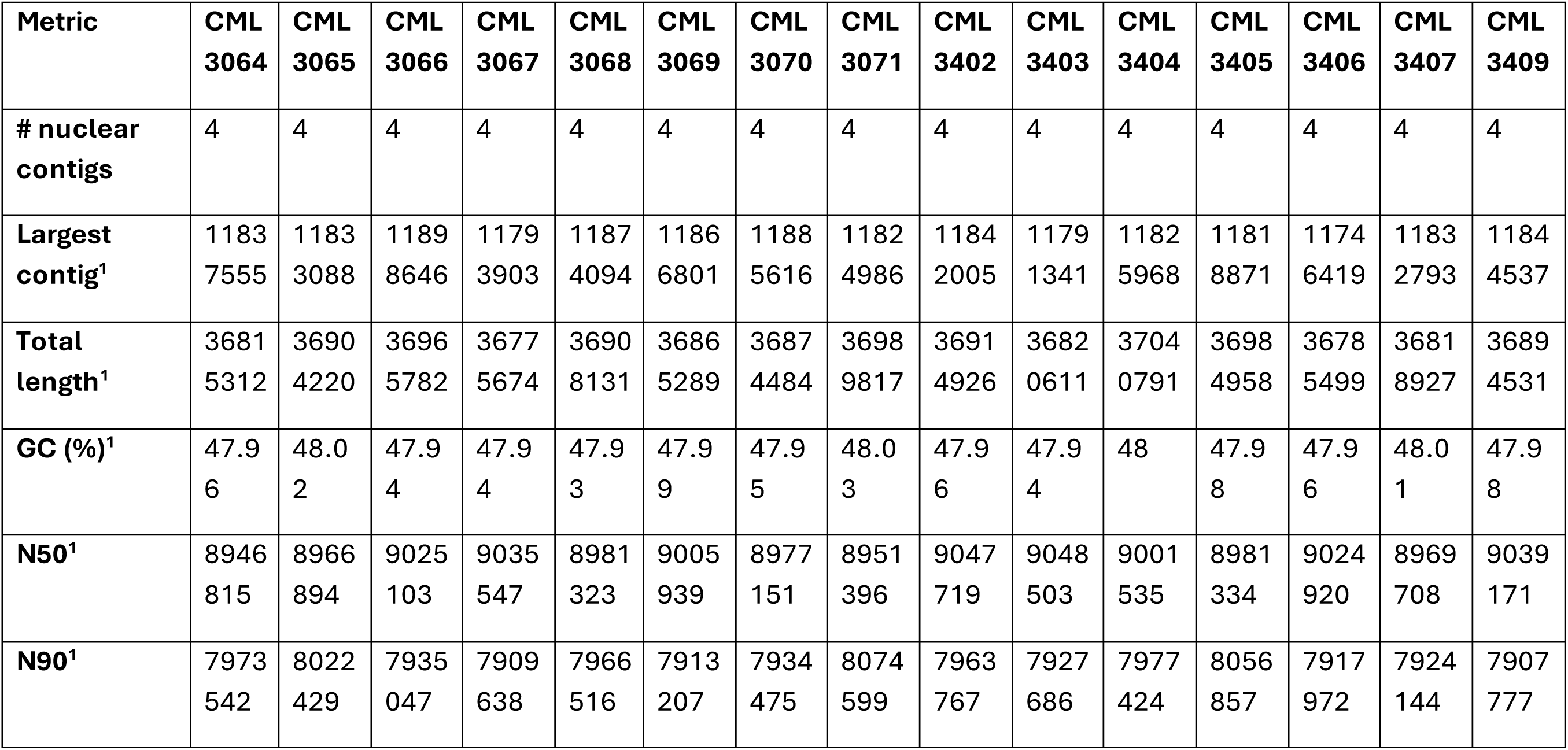
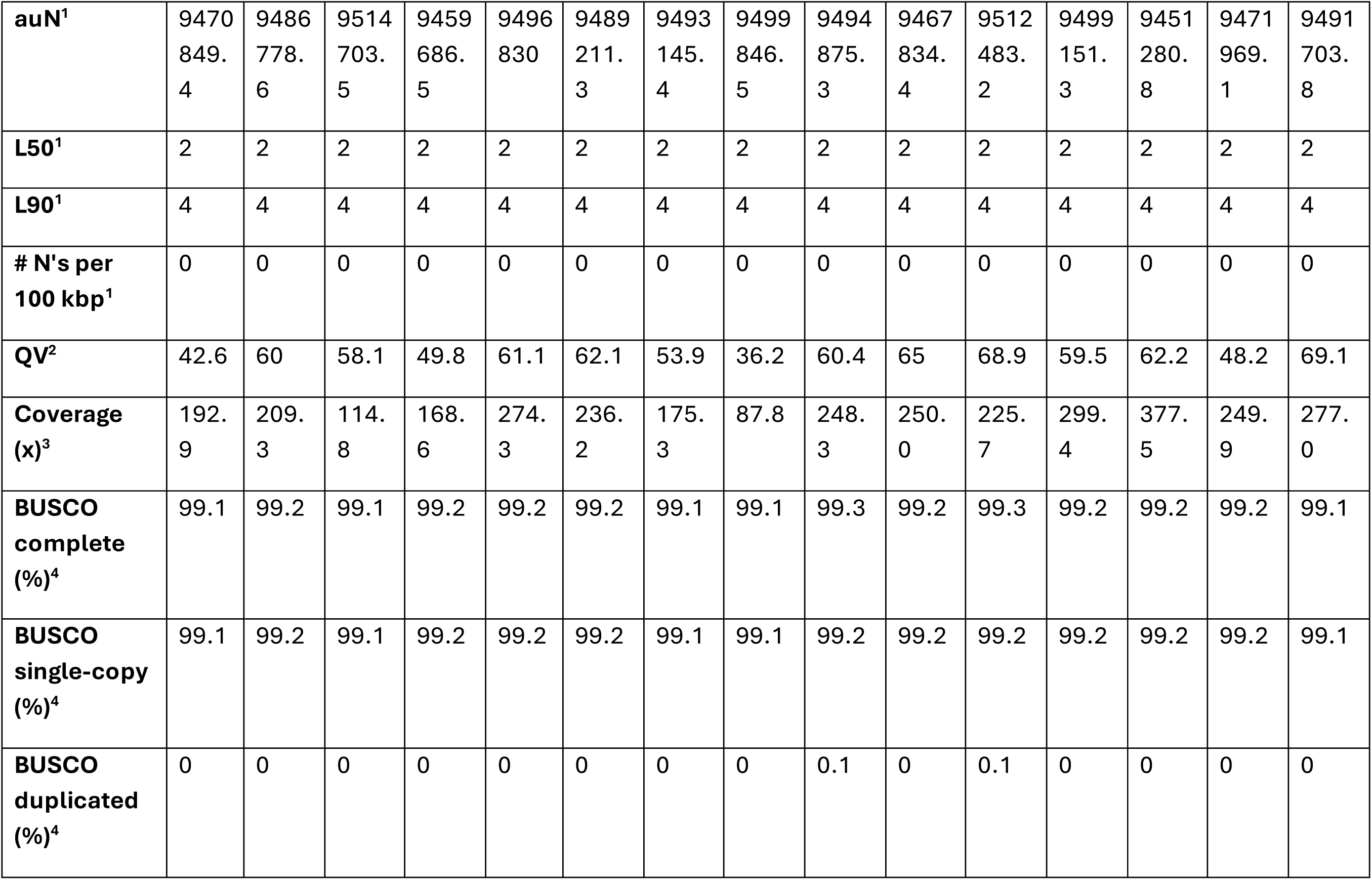
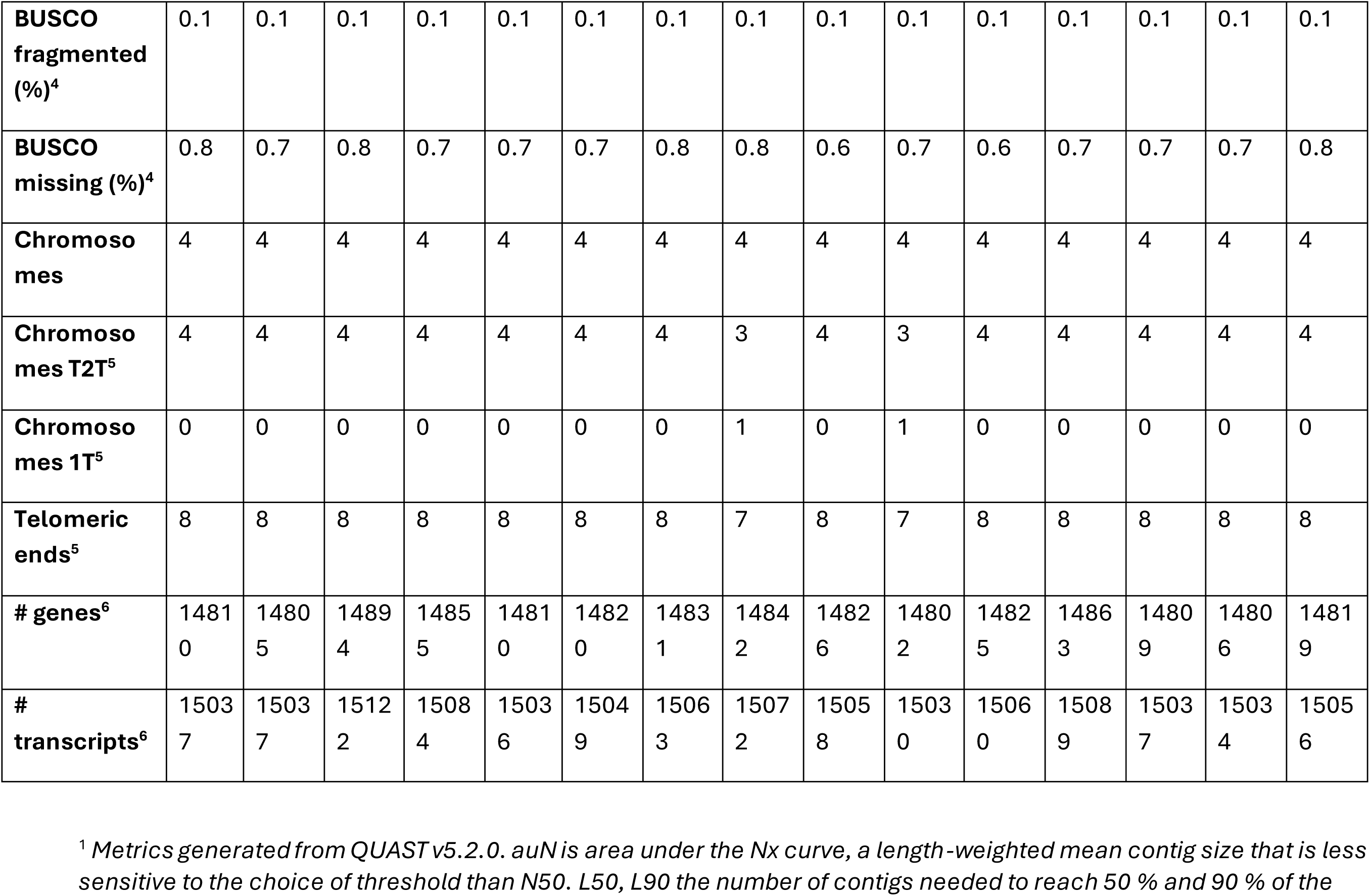
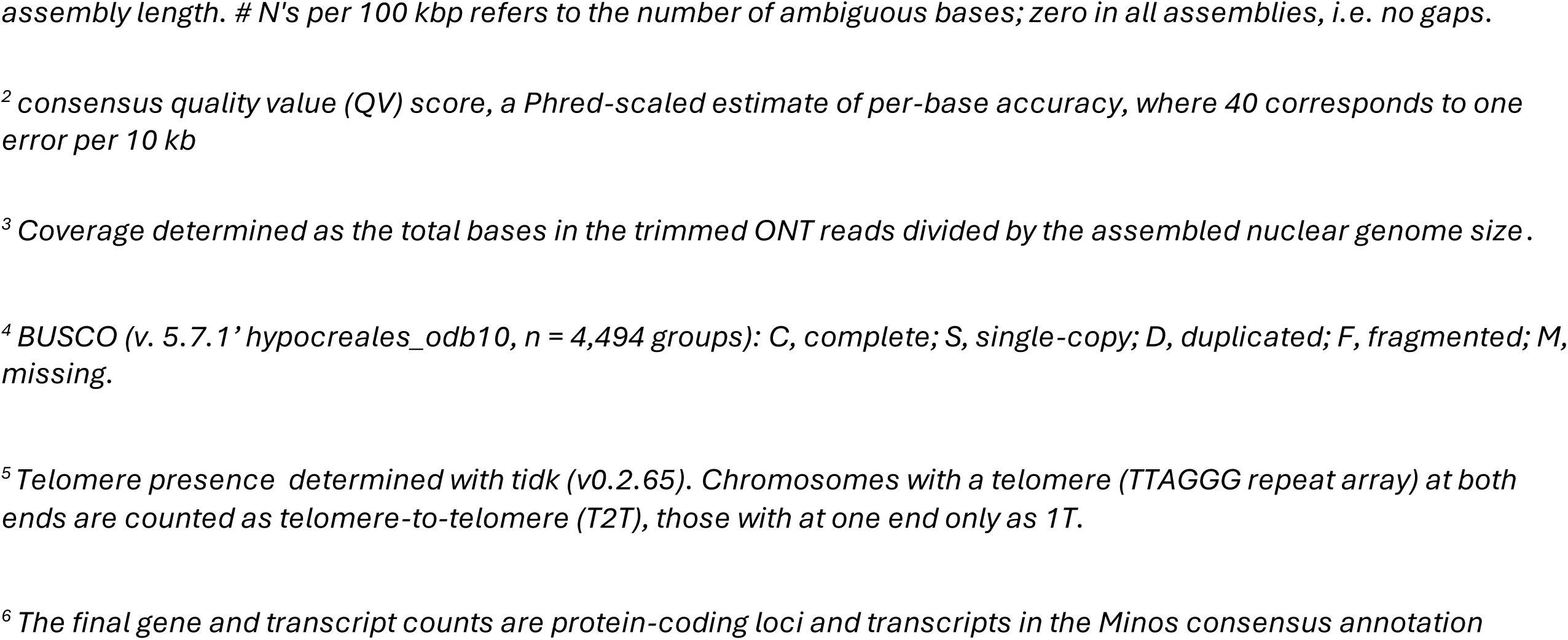
Assembly, completeness and annotation statistics for the fifteen Brazilian F. graminearum genomes. Summary of assembly metrics. Lengths are in base pairs, and auN is the area under the Nx curve. Gene-space completeness was assessed with BUSCO against the hypocreales_odb10 lineage (n = 4,4S4). Gene and transcript counts are gene and mRNA features in the final annotation.

### Gene annotation

For all 15 Brazilian isolates and the corrected PH-1 assembly (GCA_020991245.1) (Lu et al. 2022), a species-specific repeat library was built *de novo* with RepeatModeler (v2.0.4; BuildDatabase, RepeatModeler-LTRStruct) (Flynn et al. 2020). Assemblies were then masked with RepeatMasker (v4.1.5) (Smit et al. 2013).

*F. graminearum* RNA-seq reads (Dilks et al. 2019 *in vitro* dataset; ENA: PRJEB75530) were aligned to each repeat-masked assembly with STAR (v2.7.11a) (Dobin et al. 2013). Protein-coding genes were predicted with BRAKER3 (v3.0.8) (Gabriel et al. 2024) in fungus mode (--fungus), integrating both the aligned RNA-seq BAMs and protein evidence from the *F. graminearum* CML3066 v2 proteome (Wood et al. 2020) (GenBank GCA_900073075.1). To provide alternative evidence-weighted gene models for downstream consolidation, BRAKER3 was run under complementary evidence modes (RNA-seq + protein, RNA-seq only, protein only). *F. graminearum* PH-1 annotations (King et al. 2017; Lu et al. 2022) were projected onto each masked T2T assembly with Liftoff (v1.6.3) (Shumate and Salzberg 2021).

For each isolate, the BRAKER3 predictions and Liftoff-derived models were consolidated into a single non-redundant high confidence consensus gene set with Minos (v1.8.0) (https://github.com/EI-CoreBioinformatics/minos) (Venturini et al. 2018). Consensus gene coordinates were standardised and start/stop codons validated with GAG (v2.0.1) (Geib et al. 2018), producing the final per-isolate annotation.

### Orthology analysis and pangenome assignment

Orthologous groups were inferred with FastOMA (v0.3.4) (Majidian et al. 2025), using the Minos consensus proteomes of the 15 CML isolates together with the PH-1 reannotation. Proteins were placed into hierarchical orthologous groups (HOGs) against the OMAmer LUCA database (LUCA.h5), yielding root-level HOGs (RootHOGs), genes descended from a single ancestral gene, across all proteomes. The RootHOGs.tsv output was parsed into a gene-family presence/absence matrix, with isolate-specific singletons added, and gene families classified as core (present in all genomes) or accessory (present in a subset) for pangenome analysis.

### Functional annotation

Predicted proteins were functionally annotated by combining: eggNOG-mapper (v.2.1.9) (Cantalapiedra et al. 2021) (orthology, GO terms, KEGG, COG, preferred gene names), InterProScan (v.5.63) (Jones et al. 2014) (protein domains/families), and AHRD (v3.3.3) (human-readable descriptions) (https://github.com/groupschoof/AHRD). Carbohydrate-active enzymes were identified with dbCAN v3.0.1 (Zheng et al. 2023), and secondary-metabolite gene clusters with antiSMASH (v8.0.4) (Blin et al. 2025). Secreted proteins and candidate effectors were predicted using a pipeline adapted from Hill, Grey, et al. 2025. In brief, proteins were kept that carried a signal peptide according to SignalP v3, v4 and v6 (Bendtsen et al. 2004; Petersen et al. 2011; Teufel et al. 2022). Evidence was then integrated with TargetP v2 (Almagro Armenteros et al. 2019), DeepSig v.1.2.5 (Savojardo et al. 2018) and Phobius v.1.01 (Käll et al. 2004) to determine the presence of a secretion signal. This set was then filtered to remove those with more than one transmembrane domain (determined with TMHMM v.2.0c (Krogh et al. 2001) and Phobius v.1.01 (Käll et al. 2004)) or an endoplasmic-reticulum retention motif (predicted with ps_scan v.1.86 (Gattiker et al. 2002)), and finally required to be assigned an extracellular or cytoplasmic localisation by DeepLoc2 (v2.1) (Thumuluri et al. 2022). Candidate effectors were predicted from this secretome with EffectorP v1, v2 and v3 (Sperschneider et al. 2016; Sperschneider et al. 2018; Sperschneider and Dodds 2022), retaining proteins classified as effectors by at least two of the three. Pathogenicity- and virulence associated genes were mapped to the Pathogens Host Interactions (PHI-base) database (v4.18) (Urban et al. 2022; Urban et al. 2025). PHI-base annotations were assigned to PH-1 and propagated to the other genomes through the FastOMA orthogroups. Transposable elements were annotated *de novo* in each genome with Earl Grey v7.3.0 (-r ascomycota) (Baril et al. 2024).

The antiSMASH annotated regions were mapped onto the published *F. graminearum* and *venenatum* cluster catalogue (C01–C75) (Sieber et al. 2014; King et al. 2018) by orthology, using the FastOMA orthogroups to link each gene to its PH-1 counterpart. Because a region is a merged neighbourhood and may contain more than one pathway, a published cluster was counted as present only where at least two of its genes were represented. Regions were named Region_CN, or Region_CN/N where two published clusters were merged. Individual clusters were resolved at the antiSMASH proto_core, the backbone gene triggering each rule call, with genes assigned by the surrounding protocluster span.

Clusters were compared against *Fusarium venenatum*, *Fusarium verticillioides*, *Fusarium subglutinans* and *Fusarium sporotrichioides*, each annotated with antiSMASH under the same settings. Core biosynthetic genes were searched against each comparator proteome with blastp (E ≤ 1 × 10⁻⁵) (Johnson et al. 2008), accepting a homologue at ≥50% identity over ≥50% of the query length, and when present in an antiSMASH region in that species.

### Phylogenetic analysis

A species tree was inferred from single-copy orthologues shared across the 15 CML isolates, including the reannotated PH-1, and *Fusarium culmorum* (GCA_900074845.1) (Urban et al. 2016) (outgroup). From the FastOMA orthologous groups, groups present as exactly one copy in all 17 taxa were retained. Each single-copy group was aligned with MAFFT (v7.505) (Katoh and Standley 2013) and trimmed with trimAl (Capella-Gutiérrez et al. 2009). A random subsample of 1,300 trimmed alignments was concatenated into an amino-acid supermatrix with AMAS (Borowiec 2016). A maximum-likelihood phylogeny was estimated with RAxML-NG (v1.2.2) (Kozlov et al. 2019) under the JTT+I+G4 model, with 1,000 bootstrap replicates (autoMRE) and bootstrap convergence confirmed. The tree was rooted on *F. culmorum*.

### Synteny analysis

Synteny among the 15 CML isolates and PH-1 was analysed with GENESPACE v1.3.1 (Lovell et al. 2022), using the Minos gene models (GFF3 and corresponding proteomes) as input. Within GENESPACE, orthogroups were inferred with OrthoFinder (Emms and Kelly 2019) (DIAMOND v2.1.8 for all-vs-all protein search) and collinear syntenic blocks identified with MCScanX (v1.0.0) (Wang et al. 2012). Syntenic relationships were visualised as riparian plots, with genomes ordered according to the maximum-likelihood phylogeny.

## Results

### Phenotypic characterisation

Fifteen *Fusarium graminearum* isolates were collected from wheat infecting populations in southern Brazil. Aggressiveness was assessed by point inoculation of wheat cv. Bobwhite, scoring infected spikelets at 15 dpi. Isolates were classified as High where more than six spikelets became infected, Medium at three to six, and Low based on disease assessment criteria in Walkowiak et al., 2016.

The Brazilian isolates spanned the full range of phenotypic classification. Nine were classified High, alongside the PH-1 reference; two were Medium (CML_3071, CML_3068); and three were Low (CML_3404, CML_3069, CML_3067). The classes are well separated, with mean infection of 73.0% across the High isolates, 25.0% across the Medium and 7.9% across the Low isolates. All 15 isolates carry the 15-ADON trichothecene genotype, so this variation in aggressiveness is not attributable to chemotype. To determine whether these differences in aggressiveness are accompanied by differences in gene content, the genomes were sequenced and genetic content was compared at the level of orthologous gene families.

### Genome assembly and quality assessment

The fifteen Brazilian *F. graminearum* isolates were sequenced on the Oxford Nanopore platform to a median depth of 236× (Table 1). Assemblies were generated with hifiasm from either raw or HERRO-corrected reads, and contigs were ordered into chromosomes by alignment against the CML3066 v2 (Wood et al. 2020) reference. All fifteen isolates assembled into four nuclear chromosomes and one mitochondrial genome, consistent with the established *F. graminearum* karyotype (King et al. 2017; Lu et al. 2022), with total assembly sizes of 36.78–37.04 Mb (median 36.89 Mb) and GC content of 47.9–48.0%, consistent with previous assemblies (Gardiner et al. 2014; Walkowiak et al. 2016; King et al. 2017; Wood et al. 2020). Contig N50 values ranged from 8.95 to 9.05 Mb. Finally, the median Quality Variance (QV) score of the assemblies was 60.0 (Table 1).

Assembly completeness was assessed both by gene content and by telomere resolution. BUSCO analysis against the hypocreales_odb10 dataset recovered 99.1–99.3% of expected single-copy orthologues, with 0.0–0.1% duplicated and 0.6–0.8% missing (Table 1). Telomeric repeat arrays were identified with tidk at 118 of the 120 chromosome ends across the fifteen isolates, and 58 of the 60 nuclear chromosomes being resolved telomere-to-telomere. The two unresolved ends were distributed across two isolates (CML_3071 and CML_3403) (Table 1, Supplementary File 1). Mitochondrial genomes were assembled for all isolates, of which two (CML_3068 and CML_3403) were recovered as complete circular sequences.

### Comparative annotation of Brazilian isolates and PH-1

To allow gene content to be compared without the confounding effect of differing annotation methods (Hill et al. 2025), the PH-1 reference genome was re-annotated with the identical pipeline applied to the Brazilian isolates. Consolidating BRAKER3 *de novo* predictions and Liftoff-transferred PH-1 models with Minos yielded a median of 14,820 protein-coding genes per Brazilian isolate (range 14,802–14,894) and 15,346 for the re-annotated PH-1 assembly.

Functional annotation identified 724–745 predicted secreted proteins per isolate, of which 203–211 were classified as candidate effectors by EffectorP, and 1,058–1,083 carbohydrate-active enzymes by dbCAN. antiSMASH predicted 47–49 secondary metabolite regions per genome. The counts were closely similar across all sixteen genomes, and no isolate was an outlier in any category (Supplementary File 2).

### Phylogeny and synteny

A maximum-likelihood phylogeny inferred from 1,300 single-copy orthologues placed all fifteen Brazilian isolates in a single well-supported clade, with *F. culmorum* (GCA_900074845.1) as the outgroup (Figure 2A). PH-1 fell outside this clade as a sister lineage, consistent with the Brazilian population being distinct from the North American reference. Internal branch lengths within the Brazilian clade were short relative to the PH-1 divergence, indicating either a recent common origin or continuing gene flow within the sampled population.

**Figure 1.**
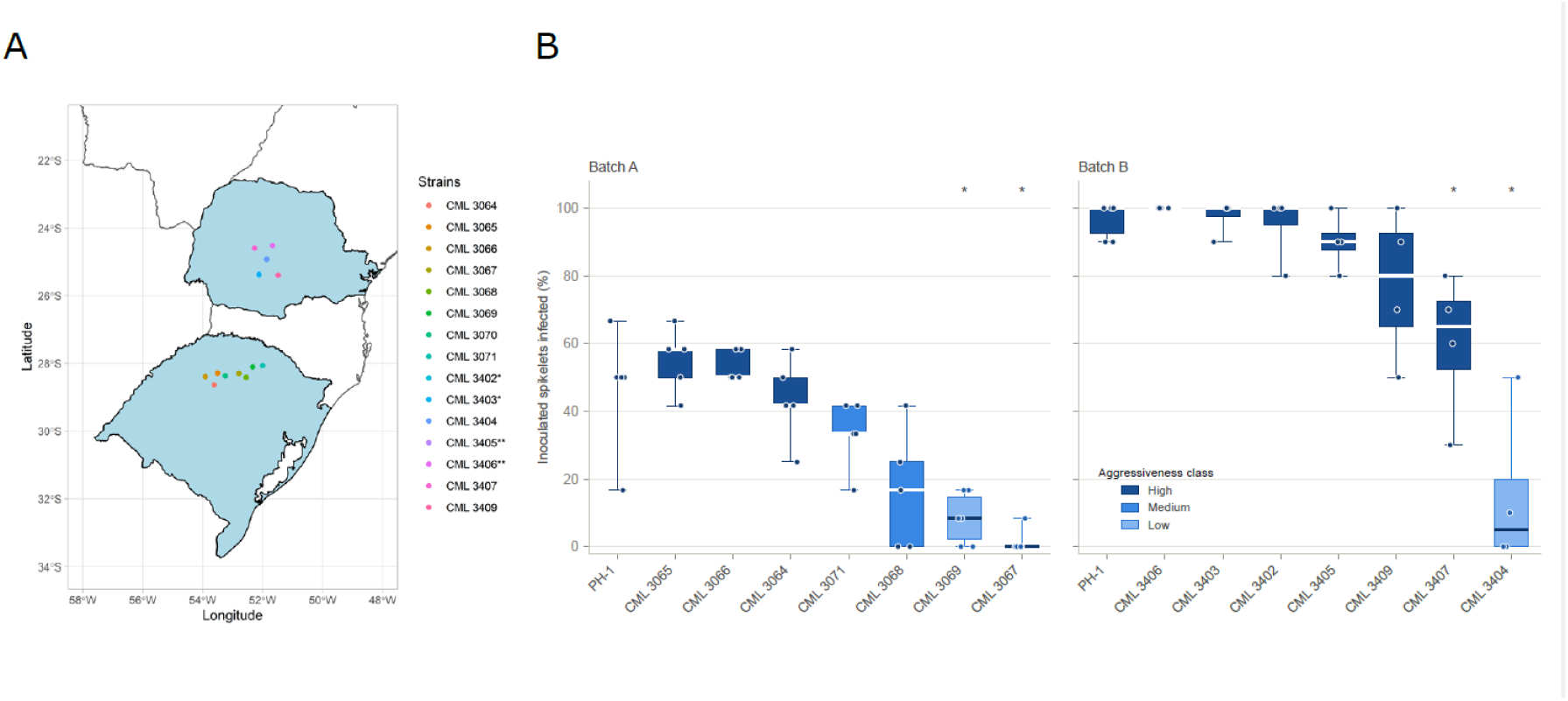
Origin and aggressiveness of the Brazilian F. graminearum isolates. A. Collection sites of the fifteen Brazilian isolates, in the states of Paraná (upper region) and Rio Grande do Sul (lower region). Each point marks the sampling location of one isolate, coloured by isolate as indicated. B. Aggressiveness on wheat cv. Bobwhite, scored as the percentage of inoculated spikelets showing symptoms at 15 days post-inoculation. Isolates were assessed in two batches; PH-1 was included in both as the reference. Fill colour denotes the aggressiveness classification. Asterisks mark isolates differing significantly from PH-1 within the same batch (two-sided Mann–Whitney U test, Benjamini–Hochberg adjusted p < 0.05).

**Figure 2.**
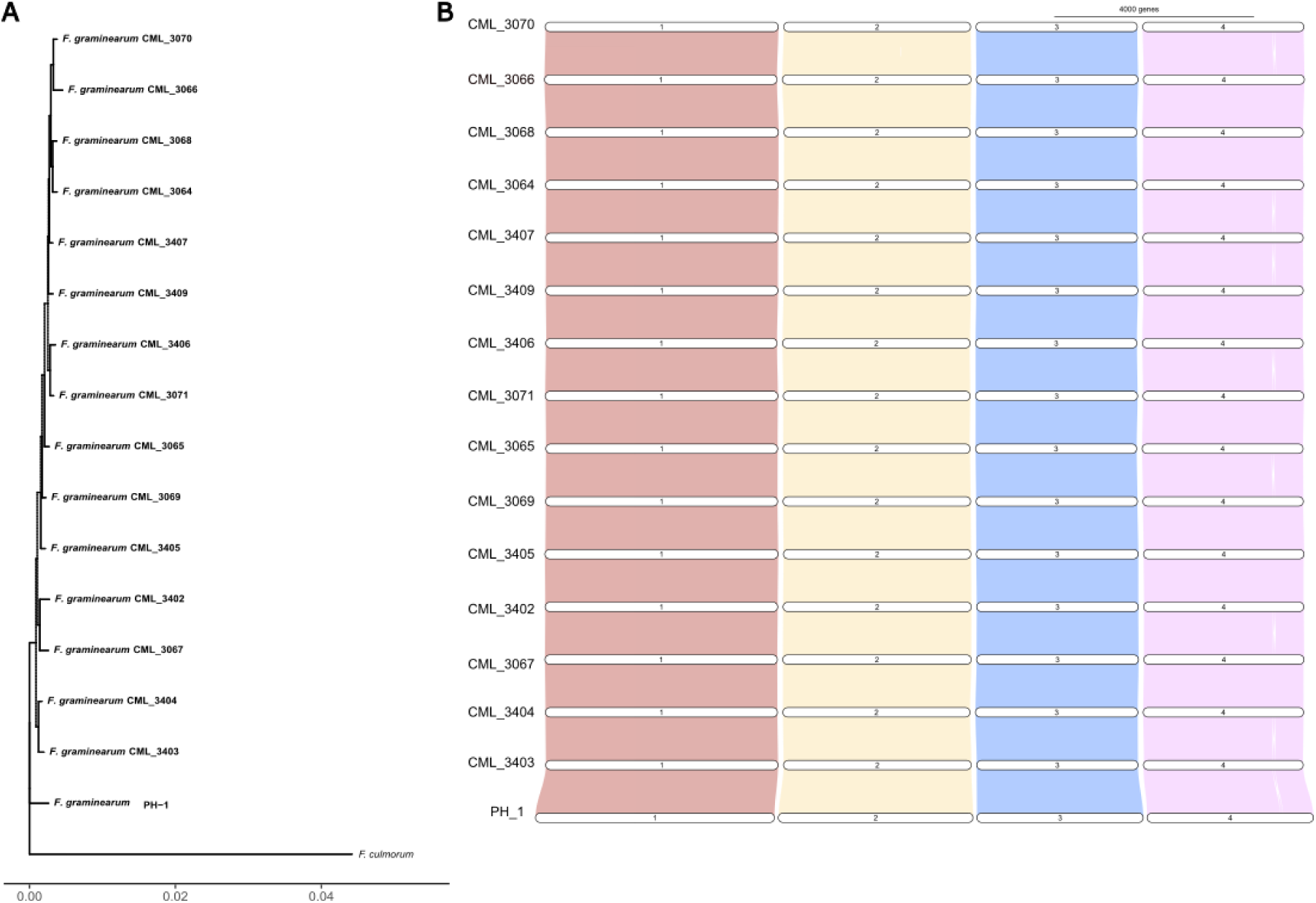
Whole-genome phylogeny and chromosome-scale synteny. A. Maximum-likelihood phylogeny from single-copy orthologues. B. Synteny plot of all 1c isolates

Whole-genome alignment showed extensive collinearity between all sixteen genomes (Figure 2B). No large-scale rearrangements were detected between the Brazilian isolates and PH-1 at the chromosome scale, and chromosome boundaries were conserved throughout. Structural variation is therefore not the dominant source of genomic difference in this population; the variation that does exist is instead presence–absence variation of gene content, which we characterise below.

### Pan-genome structure and presence–absence variation

Orthology assignment was done using FastOMA (Majidian et al. 2025) across all sixteen genomes which placed 237,763 genes into 23,673 hierarchical orthologous groups. Categories were assigned at gene level, a gene is counted present in a isolate if a representative isoform belongs to an orthogroup containing a member from that isolate. On this basis 214,847 genes (90.4%) were core, present in all sixteen genomes, 13,968 (5.9%) accessory, present in at least two genomes, and 8,948 (3.8%) isolate-specific (Figure 3). The pan-genome is therefore predominantly core.

**Figure 3.**
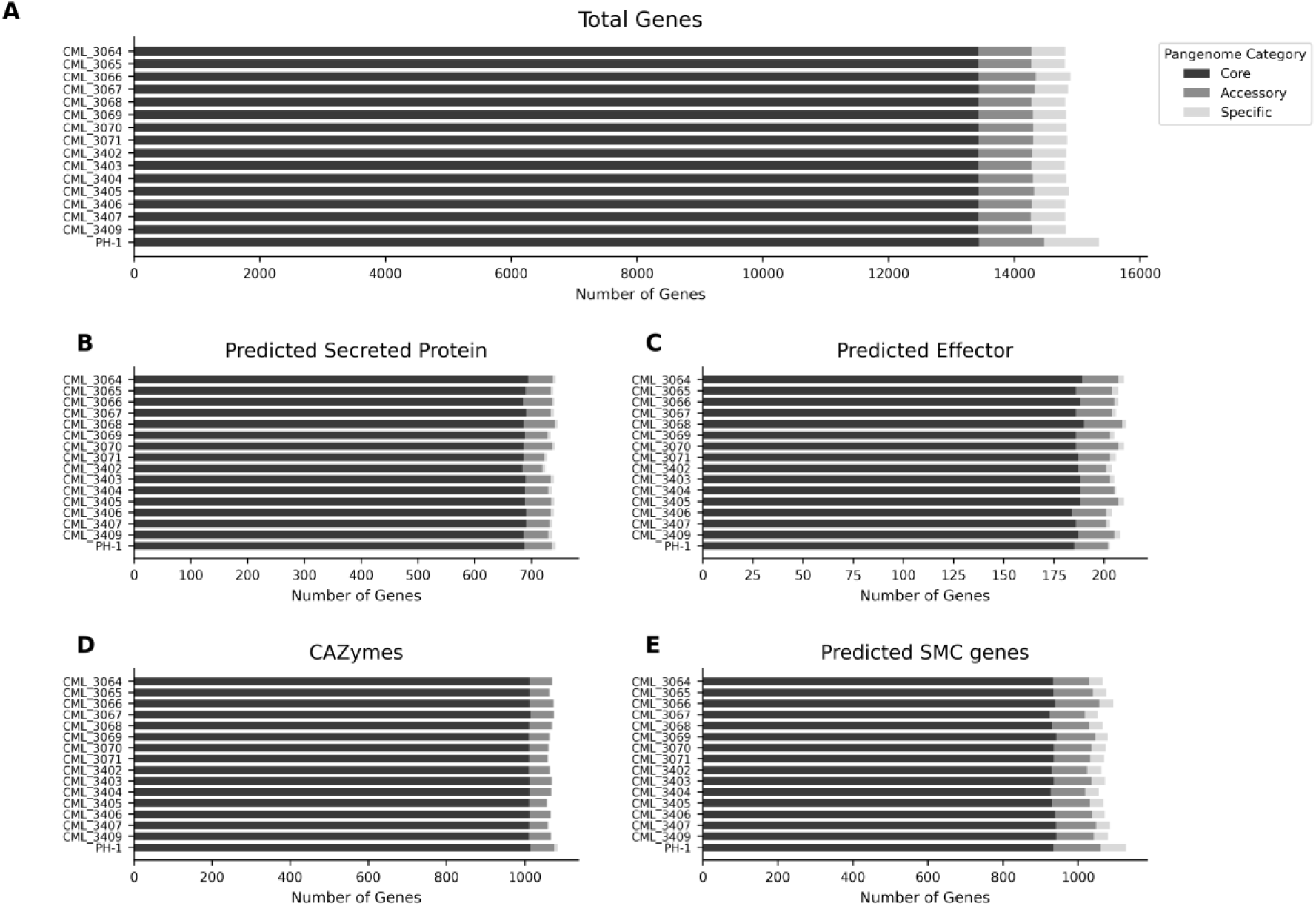
Pan-genome structure. A. Total predicted gene counts per isolate, partitioned by pangenome category (B–E) Number of genes per isolate with functional annotations for: B. predicted secreted proteins, C. predicted effector proteins, D. CAZymes (Carbohydrate-Active enZymes), and E. predicted secondary metabolite cluster (SMC) genes identified by antiSMASH. In all panels, bars are coloured by pangenome category as in A.

Gene counts were closely consistent across the Brazilian isolates (14,802–14,894 genes; 526–556 isolate-specific and 836–909 accessory each) (Figure 3–4, Supplementary File 2). PH-1 carried more of both, 1,034 accessory and 874 isolate-specific genes. However, this most likely reflects annotation bias as the transcript evidence used for gene prediction derived from PH-1. Recovering the full isolate-specific complement of the Brazilian isolates would require RNA-seq evidence from the isolates themselves. However, as the annotation bias affects all fifteen Brazilian genomes equally, comparisons among core and accessory gene content with PH-1 remain unaffected.

**Figure 4.**
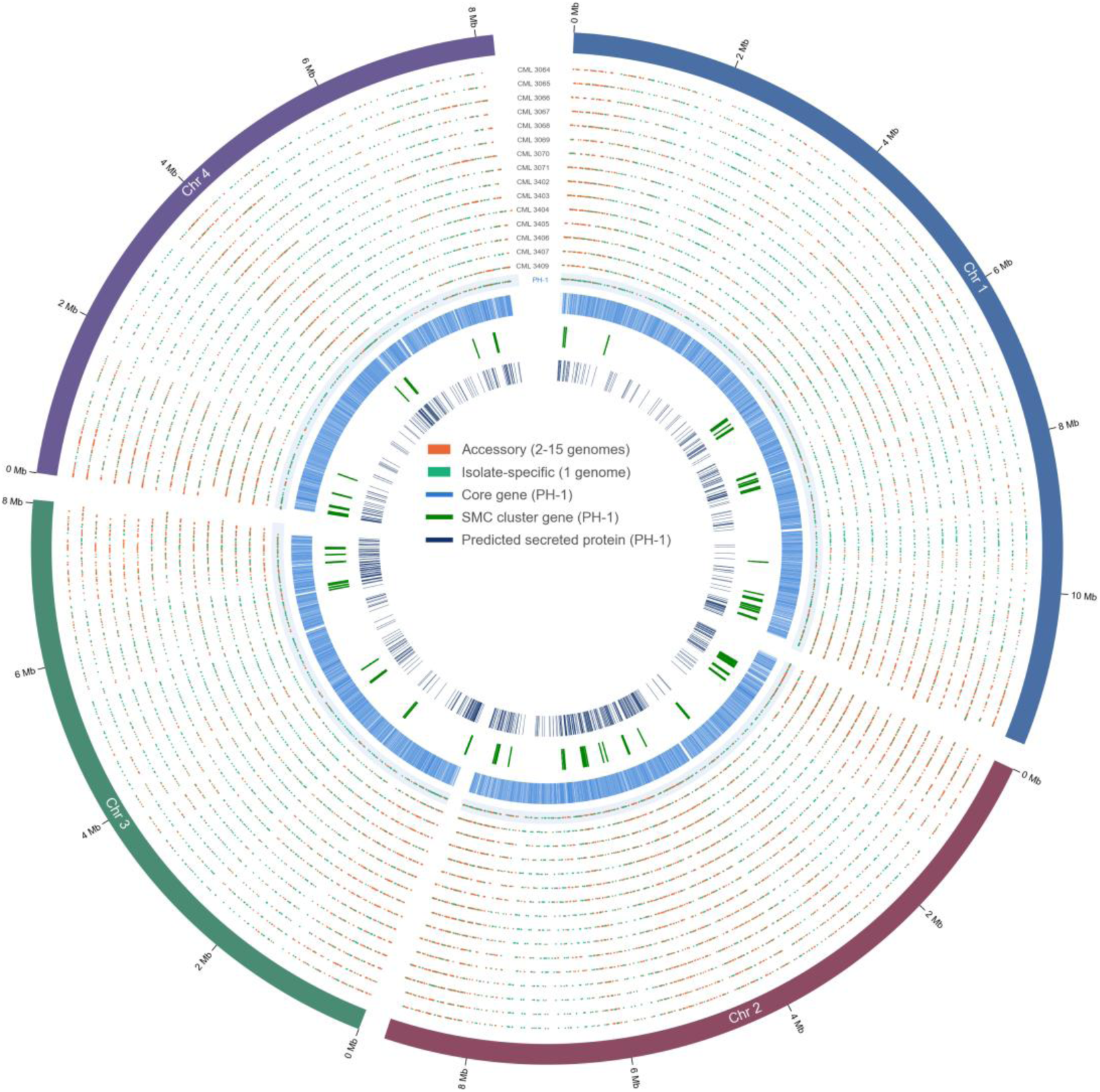
Distribution of accessory and isolate-specific genes across the *F. graminearum* Brazilian pangenome. The four F. graminearum chromosomes are drawn to scale as sectors, with positions in Mb. Each of the 15 Brazilian genomes are represented as inner rows, ordered CML_30c4 to CML_340S. PH-1 is represented at the innermost on a shaded band. Points mark accessory genes (orange, present in two to fifteen genomes) and isolate-specific genes (teal, present in one). The three innermost tracks show, for PH-1, the positions of core genes (blue), secondary metabolite cluster genes (green) and predicted secreted proteins (deep blue). The isolate specific subset is provided in Supplementary File 3.

Presence–absence variation was tabulated per gene from the same orthogroup assignment, recording this for each gene of each genome. Each gene was additionally given a synteny score that conforms to the data-format and quality-control recommendations of the ELIXIR E-PAN pan-genome framework (Heuermann et al. 2025). Of the 228,815 genes with a scoreable neighbourhood, 98.1% scored above 0.8, consistent with the collinearity seen in the whole-genome alignments. The complete pairwise table is available from the data repository (see Data Availability). Adopting a community-standard format allows this matrix to be integrated directly with other pan-genome datasets, so that presence–absence calls made here can be compared with those from other species and other studies rather than remaining a stand-alone table

### Secondary metabolite clusters

Secondary metabolite gene clusters were predicted with antiSMASH 8.0.4 in all fifteen Brazilian isolates and in the re-annotated PH-1 reference, and the predictions were mapped onto the published *F. graminearum* PH-1 and *Fusarium venenatum* clusters (C01–C75) (Sieber et al. 2014; King et al. 2018) by orthology, using the FastOMA orthogroups to link each gene to its PH-1 counterpart. antiSMASH defines a cluster region as a merged neighbourhood, which can merge neighbouring protocluster regions into a single cluster. The antismash cluster regions therefore do not correspond one-to-one with the previously published clusters, specifically 3 of 51 antismash regions contained two previously published clusters rather than one. To carry the established nomenclature forward, antismash cluster regions were named Region_CN, or Region_CN/N where a new cluster now merged two previously published clusters. Individual clusters were resolved at the level of the antiSMASH proto_core, the backbone biosynthetic gene that triggers each rule call, and clusters with no published counterpart were grouped across genomes by orthogroup content so that one locus carries one name in every isolate. These were named as region_C76 onward, naming new cluster cores C76–C88 to continue from the naming in King et al., 2018.

A total of 53 distinct clusters were identified across the sixteen genomes, including 13 new ones which did not map to previously identified clusters. 49 of the 53 clusters were present in all sixteen genomes, including the trichothecene, aurofusarin, zearalenone, fusarielin and fusaoctaxin A clusters, with four cluster cores varying in presence (Table 2).

**Table 2.** Secondary metabolite clusters with a variable core gene across the sixteen genomes. Presence is scored on the backbone biosynthetic gene, the antiSMASH core that triggers each rule call. These are all four of the 53 cluster cores that are not detected in every genome. Genomes present gives the number of genomes in which the core gene was predicted; the product class is that of the protocluster the core heads, and the metabolite that of the closest MIBiG reference cluster where one was assigned.

| Cluster core | Core enzyme | Product class | Metabolite | Genomes present | Isolates carrying the core |
| --- | --- | --- | --- | --- | --- |
| C47 | NRPS7; PKS6 | NRPS, T1PKS | fusaristatin A | 14/16 | all except CML_3067 and CML_3404 |
| C01 | UbiA<br>prenyltransferase | terpene | — | 6/16 | CML_3065,<br>CML_3066,<br>CML_3068,<br>CML_3069,<br>CML_3407,<br>CML_3409 |
| C88 | PKS52 | T1PKS | alternapyrone-<br>like | 5/16 | CML_3067,<br>CML_3069,<br>CML_3403,<br>CML_3406,<br>CML_3409 |
| C81 | Geranylgeranyl<br>pyrophosphate<br>synthase | terpene | Gibberellin-<br>like | 3/16 | CML_3064,<br>CML_3066,<br>CML_3068 |

The core trichothecene cluster (C23) is present in all sixteen genomes, but its gene content varies. Namely, TRI7 is absent from seven Brazilian isolates (CML_3065, CML_3066, CML_3070, CML_3071, CML_3402, CML_3405 and CML_3409). TRI7 acetylates C-4 and its loss is the established determinant of the 15-ADON chemotype, so this variation is consistent with the chemotype observed among the Brazilian isolates.

The newly identified C88 is present in five isolates, CML_3067, CML_3069, CML_3403, CML_3406 and CML_3409, and absent from the other eleven. It is defined by a type I polyketide synthase corresponding to PKS52 (Hansen et al. 2015), which is absent from the PH-1 reference re-annotated here. Its genes share 51–71% amino acid identity with those of the alternapyrone pathway of the potato pathogen *Alternaria solani*, in which the iterative polyketide synthase PKSN produces the octa-methylated decaketide alternapyrone (Fujii et al. 2005), a compound of unknown biological function. Its distribution does not, however, follow aggressiveness, as the five carriers comprise three isolates classified High and two classified Low. It is therefore unclear how the metabolite might contribute to *F. graminearum* life cycle. As the product of C88 is unidentified, and a metabolite of this class may act conditionally, under host genotypes, tissues or environmental conditions not captured by point inoculation of a single cultivar, or in combination with other factors that differ between these isolates. Establishing whether C88 is functionally relevant will require expression and metabolite data from the carrier isolates rather than presence–absence alone.

To identify which clusters are restricted *to F. graminearum*, the sixteen genomes were compared against four species spanning different hosts and lifestyles*: F. venenatum*, which is non-pathogenic towards wheat spikes (King et al. 2018), and *F. verticillioides, F. subglutinans* and *F. sporotrichioides*, which infect maize and other cereals. Four of the clusters present in all sixteen *F. graminearum* genomes have no core biosynthetic gene homologue in the comparator species. These clusters are C02, a fusarielin H cluster (C60), C61, and the fusaoctaxin A cluster (C64). The Fusaoctaxin A is an established cluster with the characterised core gene *Nrps9* known to be wheat virulence factor required for cell-to-cell invasion, and its restriction to *F. graminearum* among these species is consistent with a role in the wheat-adapted lifestyle (Jia et al. 2019). The other three clusters (C02, C60 and C61) do not have their core genes characterised but could be similarly important for virulence. Of the newly identified clusters 8 out of 13 have a counterpart *in F. venenatum*, their backbone gene being orthologous to a gene that antiSMASH also placed within a predicted cluster in that species, and in every case the two clusters were assigned the same product class independently (Supplementary File 4).

### PHI-base and pathogenicity and virulence associated genes

Each Brazilian isolate carries 1,231–1,242 PHI-base-matched genes, 1,211–1,216 of them core and only 17–29 accessory, so pathogenicity and virulence associated variation is confined to a small accessory set. Three genes with a reported reduced-virulence phenotype vary across the panel. These are FGSG_03243 (12/16), FGSG_03624 (11/16), and the orphan secreted protein Osp44 (7/16). Notably, FGSG_03243 and Osp44 are both absent from the low aggressive isolate CML_3404. FGSG_03243, FGSG_03624 and Osp44 are predicted secreted proteins, with Osp44 being a characterised effector (Jiang et al. 2020). This therefore suggests a possible association of varying effector repertoires with virulence.

### Transposable elements

Transposable elements were annotated *de novo* in each genome with EarlGrey. Transposable element content was uniform across the population with 2,572–2,741 annotated copies per genome. Ninety-four families had copies in all sixteen genomes. Five families were detected in only one genome, but each was represented by a single copy and all derived from the curated library of other fungal species rather than from *de novo F. graminearum* families, so none constitutes evidence of isolate-specific repeat content. Length-based completeness is a proxy for structural intactness and not for transposition competence, since open reading frames and terminal repeats were not assessed. That transposable element content is uniform across the population, indicates an absence of recent lineage-specific proliferation. This is consistent with the structural conservation of these genomes and with repeat-induced point mutation, which is active in *F. graminearum* (Cuomo et al. 2007), constraining repeat expansion.

## Discussion

The fifteen assemblies reported here, constitute a high resolution multi-genome resource assembled for *F. graminearum* that is also the first to represent the diversity in the Brazilian populations.

The 90.4% core reported here is much closer to the 12% accessory fraction described across ten genomes of the *F. graminearum* species complex (Walkowiak et al. 2016) than to the 56% accessory reported within strains of F. graminearum alone (Alouane et al. 2021). Since the former comparison spans three species and the latter one, it suggests that the more variable pangenome may have been due to variable quality in assemblies and annotations. Here, with 58 of 60 nuclear chromosomes resolved telomere-to-telomere and consistent annotations across all isolates, every scored absence is likely an absence from assembled sequence rather than from an assembly or annotation gap.

The genomic uniformity of this population is striking given the phenotypic range it spans. The isolates differ more than nine-fold in mean spikelet infection, from 7.9% to 73.0%, yet share the same karyotype and chemotype, carry near-identical transposable element content with no isolate-specific family expansions, and vary minimally in gene count. One explanation is that *F. graminearum* has comparatively little to gain from genomic diversification because its infection strategy is so tightly coupled to mycotoxin production, notably deoxynivalenol. Trichothecene biosynthesis is required for the fungus to grow through the rachis node and colonise the spike beyond the inoculated spikelet (Proctor et al. 1995; Jansen et al. 2005; Armer, Urban, et al. 2024), and DON acts by inhibiting eukaryotic translation, a mechanism that is largely indiscriminate with respect to host genotype. A pathogen whose principal determinant of colonisation is a conserved, chromosomally integrated toxin with a species non-specific mode of action is not subject to the gene-for-gene recognition dynamics that drive rapid effector turnover. Consistent with this, species of the *Fusarium sambucinum* clade have among the lowest chromosome numbers in the genus and few to no accessory chromosomes (Armer, Kroll, et al. 2024), while the two-speed genome of *F. graminearum* is expressed as an intragenomic subtelomeric compartment rather than as dispensable chromosomes (Wang et al. 2017). Perhaps a strong reliance on mycotoxins to invade the highly reinforced grain-bearing tissues of cereals has permitted the loss of other virulence components from the genome, leaving a compact and largely invariant gene set. If gene content is largely fixed, the aggressiveness differences documented here must be substantially regulatory, or must reside in the small accessory fraction. However, distinguishing these explanations will require expression data from and experimental charactersation of the Brazilian isolates themselves.

In a landscape where only around one-fifth of publicly available fungal genome assemblies are long-read and just a quarter carry gene annotations (Kroll et al. 2026), fifteen near-T2T assemblies with uniform functional annotation provide a high quality resource. Combined with a gene-level presence-absence table with synteny scores conforming to the ELIXIR E-PAN recommendations (Heuermann et al. 2025), and matched aggressiveness phenotypes, this therefore acts as a foundational baseline for FHB research.

## Supplementary

**Supplementary Table 1.** Summary of *F. graminearum* isolates used in this study. This table lists the PH-1 reference and the fifteen Brazilian isolates, summarising for each the trichothecene chemotype, the state and country of collection, the year of isolation, and the aggressiveness classification on wheat. Disease assessment as described in Walkowiak et al. 2016-High = more than six spikelets infected, Medium = three to six spikelets infected, Low = infection on spikelets adjacent to inoculation site and NI = No information. CML: Coleção Micológica de Lavras (Brazil) – Wood et al. (2020).

| Code | Species | Tricothecene<br>genotype | Geographic<br>origin | Year of<br>isolation | Aggressiveness |
| --- | --- | --- | --- | --- | --- |
| PH-1 | *F. graminearum* | 15-ADON | USA | 1996 | High |
| CML 3064 | *F. graminearum* | 15-ADON | Rio Grande<br>do Sul<br>Brazil | 2007 | High |
| CML 3065 | *F. graminearum* | 15-ADON | Rio Grande<br>do Sul<br>Brazil | 2009 | High |
| CML 3066 | *F. graminearum* | 15-ADON | Rio Grande<br>do Sul<br>Brazil | 2010 | High |
| CML 3067 | *F. graminearum* | 15-ADON | Rio Grande<br>do Sul<br>Brazil | 2010 | Low |
| CML 3068 | *F. graminearum* | 15-ADON | Rio Grande<br>do Sul<br>Brazil | 2007 | Medium |
| CML 3069 | *F. graminearum* | 15-ADON | Rio Grande<br>do Sul<br>Brazil | 2010 | Low |
| CML 3070 | *F. graminearum* | 15-ADON | Rio Grande<br>do Sul<br>Brazil | 2011 | NI |
| CML 3071 | *F. graminearum* | 15-ADON | Rio Grande<br>do Sul<br>Brazil | 2010 | Medium |
| CML 3402 | *F. graminearum* | 15-ADON | Parana<br>Brazil | 2011 | High |
| CML 3403 | *F. graminearum* | 15-ADON | Parana<br>Brazil | 2011 | High |
| CML 3404 | *F. graminearum* | 15-ADON | Parana<br>Brazil | 2011 | Low |
| CML 3405 | *F. graminearum* | 15-ADON | Parana<br>Brazil | 2011 | High |
| CML 3406 | *F. graminearum* | 15-ADON | Parana<br>Brazil | 2011 | High |
| CML 3407 | *F. graminearum* | 15-ADON | Parana<br>Brazil | 2011 | High |
| CML 3409 | *F. graminearum* | 15-ADON | Parana<br>Brazil | 2011 | High |

**Supplementary File 1. Telomere identification in the fifteen Brazilian assemblies.**

*Telomeric repeats were identified with tidk using the TTAGGG motif in 10 kb windows. For each chromosome the table gives its length and the repeat count in the terminal window at each end. An end was scored as telomeric where its terminal window contained at least five repeats; terminal windows carried a median of 32 repeats against a median of 2 in interior windows. Chromosomes with an array at both ends are recorded as telomere-to-telomere*.

**Supplementary File 2. Functional annotations of 15 Brazilian isolates and PH-1**

*Excel workbook containing the complete functional annotation of all 15 Brazilian isolates and the re-annotated PH-1 genome, one sheet per isolate. The first sheet defines every column, giving its source tool and a description. Each isolate sheet holds one row per predicted gene with its functional annotation (eggNOG-mapper, InterProScan, AHRD), CAZyme assignment (dbCAN), secondary metabolite cluster membership (antiSMASH), secretion and effector prediction, PHI-base annotation, orthogroup assignment (FastOMA) and pan-genome category*.

**Supplementary File 3. Isolate-specific genes of the fifteen Brazilian *F. graminearum* isolates and PH-1.**

*One sheet per genome, giving position, functional annotation, secretion and effector predictions, PHI-base match, secondary metabolite cluster membership and orthogroup for each of the genes whose orthogroup is restricted to a single genome. The first sheet defines every column*.

**Supplementary File 4. The presence and absence of the 13 newly assigned metabolite clusters.**

*For each cluster, its antiSMASH product class, backbone enzyme, the number of the sixteen genomes carrying it, and whether the orthologous locus is also predicted as a cluster in Fusarium venenatum, with that region’s identifier, product class, backbone gene and gene complement*.

## Data availability

Raw sequencing reads, nuclear and mitochondrial genome assemblies, and the annotations for the fifteen Brazilian isolates are available from NCBI under BioProject PRJNA1479928. All workflows and scripts used to generate and analyse these data are available at https://github.com/RothResearch/fg-brazilian-pangenome and archived at Zenodo (DOI: 10.5281/zenodo.22145692). Additional derived data are deposited at Zenodo, comprising the FastOMA orthogroup assignments (RootHOGs.tsv), the complete EffectorP predictions, the antiSMASH output for all sixteen genomes and for the four comparator *Fusarium* species, the presence–absence variation table, the transposable element annotations, and the per-genome functional annotation tables as CSV files. These functional annotation tables are also provided as Supplementary File 2.

## Acknowledgements

The authors would like to thank Dr Yedemon Ange B. Zoclanclounon at Rothamsted Research, Dr Rowena Hill at the Earlham Institute and Drs Dan Smith and Rob King formerly of Rothamsted Research for their valuable suggestions and insights into this project. We would also like to that Professor Ludwig Pfenning at the Federal University of Lavras, Brazil for his help with the preparation of the official paperwork that permitted the export of the 15 Brazilian *Fusarium graminearum* isolates from Brazil to the UK. We also thank Professor Mauricio Fernandes, formerly of EMBRAPA Trigo, for helping to co-design and support this long-term UK-Brazil bilateral project.

## Funding

This study was supported by funding from the Biotechnology and Biological Sciences Research Council (BBSRC, https://www.ukri.org/councils/bbsrc/), including the BBSRC-EMBRAPA Bilateral grants BB/N004493/1 ( K.H.K. and M.U.) and BB/N018095/1 (A.M.W, K.H.K. and M.U.), the Designing Future Wheat programme grant BBS/E/C/000I0250 (K.H.K., M.U., K.H.P, A.D.K) and the Delivering Sustainable Wheat programme grant (BB/X011003/1) within the work package Delivering Resilience to Biotic Stress BBS/E/RH/230001B Rothamsted Research) (K.H.K., M.U.). EK was supported by the BBSRC Core Capability Grant (BB/CCG2280/1).

## Bibliography

Almagro Armenteros JJ et al. 2019. Detecting sequence signals in targeting peptides using deep learning. Life Sci Alliance. 2(5):e201900429. 10.26508/lsa.201900429

Alouane T et al. 2021. Comparative Genomics of Eight *Fusarium graminearum* Strains with Contrasting Aggressiveness Reveals an Expanded Open Pangenome and Extended Effector Content Signatures. Int J Mol Sci. 22(12):6257. 10.3390/ijms22126257

Armer VJ, Urban M, et al. 2024. The trichothecene mycotoxin deoxynivalenol facilitates cell-to-cell invasion during wheat-tissue colonization by *Fusarium graminearum*. Mol Plant Pathol. 25(6):e13485

Armer VJ, Kroll E, et al. 2024. Navigating the *Fusarium* species complex: Host-Range Plasticity and Genome Variations. Fungal Biol. 128(8 Pt B):2439–2459. 10.1016/j.funbio.2024.07.004

Baril T, Galbraith J, Hayward A. 2024. Earl Grey: A Fully Automated User-Friendly Transposable Element Annotation and Analysis Pipeline. Mol Biol Evol. 41(4):msae068. 10.1093/molbev/msae068

Bendtsen JD, Nielsen H, von Heijne G, Brunak S. 2004. Improved prediction of signal peptides: SignalP 3.0. J Mol Biol. 340(4):783–795. 10.1016/j.jmb.2004.05.028

Blin K et al. 2025. antiSMASH 8.0: extended gene cluster detection capabilities and analyses of chemistry, enzymology, and regulation. Nucleic Acids Res. 53(W1):W32–W38. 10.1093/nar/gkaf334

Borowiec ML. 2016. AMAS: a fast tool for alignment manipulation and computing of summary statistics. PeerJ. 4:e1660. 10.7717/peerj.1660

Brown MR, Manuel Gonzalez de La Rosa P, Blaxter M. 2025. tidk: a toolkit to rapidly identify telomeric repeats from genomic datasets. Bioinformatics. 41(2):btaf049. 10.1093/bioinformatics/btaf049

Cantalapiedra CP et al. 2021. eggNOG-mapper v2: Functional Annotation, Orthology Assignments, and Domain Prediction at the Metagenomic Scale. Mol Biol Evol. 38(12):5825–5829. 10.1093/molbev/msab293

Capella-Gutiérrez S, Silla-Martínez JM, Gabaldón T. 2009. trimAl: a tool for automated alignment trimming in large-scale phylogenetic analyses. Bioinformatics. 25(15):1972–1973. 10.1093/bioinformatics/btp348

de Chaves MA et al. 2022. Fungicide Resistance in *Fusarium graminearum* Species Complex. Curr Microbiol. 79(2):62. 10.1007/s00284-021-02759-4

Chen H et al. 2023. Combined pangenomics and transcriptomics reveals core and redundant virulence processes in a rapidly evolving fungal plant pathogen. BMC Biology. 21(1):24. 10.1186/s12915-023-01520-6

Cheng H et al. 2026. Efficient near-telomere-to-telomere assembly of nanopore simplex reads. Nature. 655(8121):166–173. 10.1038/s41586-026-10105-6

Cuomo CA et al. 2007. The *Fusarium graminearum* Genome Reveals a Link Between Localized Polymorphism and Pathogen Specialization. Science. 317(5843):1400–1402. 10.1126/science.1143708

Danecek P et al. 2021. Twelve years of SAMtools and BCFtools. Gigascience. 10(2):giab008. 10.1093/gigascience/giab008

Dhakal U, Kim H-S, Toomajian C. 2024. The landscape and predicted roles of structural variants in *Fusarium graminearum* genomes. G3 (Bethesda). 14(6):jkae065. 10.1093/g3journal/jkae065

Dilks T et al. 2019. Non-canonical fungal G-protein coupled receptors promote Fusarium head blight on wheat. PLoS Pathog. 15(4):e1007666

Dobin A et al. 2013. STAR: ultrafast universal RNA-seq aligner. Bioinformatics. 29(1):15–21. 10.1093/bioinformatics/bts635

Emms DM, Kelly S. 2019. OrthoFinder: phylogenetic orthology inference for comparative genomics. Genome Biol. 20(1):238. 10.1186/s13059-019-1832-y

Flynn JM et al. 2020. RepeatModeler2 for automated genomic discovery of transposable element families. Proc Natl Acad Sci U S A. 117(17):9451–9457. 10.1073/pnas.1921046117

Fujii I et al. 2005. An iterative type I polyketide synthase PKSN catalyzes synthesis of the decaketide alternapyrone with regio-specific octa-methylation. Chem Biol. 12(12):1301–1309. 10.1016/j.chembiol.2005.09.015

Gabriel L et al. 2024. BRAKER3: Fully automated genome annotation using RNA-seq and protein evidence with GeneMark-ETP, AUGUSTUS, and TSEBRA. Genome Res. 34(5):769–777. 10.1101/gr.278090.123

Gardiner DM, Stiller J, Kazan K. 2014. Genome Sequence of *Fusarium graminearum* Isolate CS3005. Genome Announc. 2(2):e00227–14. 10.1128/genomeA.00227-14

Gattiker A, Gasteiger E, Bairoch A. 2002. ScanProsite: a reference implementation of a PROSITE scanning tool. Appl Bioinformatics. 1(2):107–108

Geib SM et al. 2018. Genome Annotation Generator: a simple tool for generating and correcting WGS annotation tables for NCBI submission. Gigascience. 7(4):1–5. 10.1093/gigascience/giy018

Gurevich A, Saveliev V, Vyahhi N, Tesler G. 2013. QUAST: quality assessment tool for genome assemblies. Bioinformatics. 29(8):1072–1075. 10.1093/bioinformatics/btt086

Hansen FT et al. 2015. An update to polyketide synthase and non-ribosomal synthetase genes and nomenclature in *Fusarium*. Fungal Genet Biol. 75:20–29. 10.1016/j.fgb.2014.12.004

Heuermann MC et al. 2025. White paper: standards for handling and analyzing plant pan-genomes. F1000Res. 14:739. 10.12688/f1000research.166538.3

Hill R, Reynolds G, Hall N, Swarbreck D. 2025. Leveraging existing data to maximise quality and consistency across gene model annotations: a *Fusarium* pan-annotation. 2025.03.12.642647 [accessed 2026 Apr 16]. https://www.biorxiv.org/content/10.1101/2025.03.12.642647v1. 10.1101/2025.03.12.642647

Jansen C et al. 2005. Infection patterns in barley and wheat spikes inoculated with wild-type and trichodiene synthase gene disrupted *Fusarium graminearum*. Proceedings of the National Academy of Sciences. 102(46):16892–16897. 10.1073/pnas.0508467102

Jia L-J et al. 2019. A linear nonribosomal octapeptide from *Fusarium graminearum* facilitates cell-to-cell invasion of wheat. Nat Commun. 10:922. 10.1038/s41467-019-08726-9

Jiang C et al. 2020. An orphan protein of *Fusarium graminearum* modulates host immunity by mediating proteasomal degradation of TaSnRK1α. Nat Commun. 11(1):4382. 10.1038/s41467-020-18240-y

Johnson M et al. 2008. NCBI BLAST: a better web interface. Nucleic Acids Research. 36(suppl_2):W5–W9. 10.1093/nar/gkn201

Jones P et al. 2014. InterProScan 5: genome-scale protein function classification. Bioinformatics. 30(9):1236–1240. 10.1093/bioinformatics/btu031

Käll L, Krogh A, Sonnhammer ELL. 2004. A combined transmembrane topology and signal peptide prediction method. J Mol Biol. 338(5):1027–1036. 10.1016/j.jmb.2004.03.016

Katoh K, Standley DM. 2013. MAFFT multiple sequence alignment software version 7: improvements in performance and usability. Mol Biol Evol. 30(4):772–780. 10.1093/molbev/mst010

Kelly AC, Ward TJ. 2018. Population genomics of *Fusarium graminearum* reveals signatures of divergent evolution within a major cereal pathogen. PLOS ONE. 13(3):e0194616. 10.1371/journal.pone.0194616

King R et al. 2015. The completed genome sequence of the pathogenic ascomycete fungus *Fusarium graminearum*. BMC Genomics. 16(1):544. 10.1186/s12864-015-1756-1

King R, Brown NA, Urban M, Hammond-Kosack KE. 2018. Inter-genome comparison of the Quorn fungus *Fusarium venenatum* and the closely related plant infecting pathogen *Fusarium graminearum*. BMC Genomics. 19(1):269. 10.1186/s12864-018-4612-2

King R, Urban M, Hammond-Kosack KE. 2017. Annotation of *Fusarium graminearum* (PH-1) Version 5.0. Genome Announc. 5(2):e01479–16. 10.1128/genomeA.01479-16

Kozlov AM et al. 2019. RAxML-NG: a fast, scalable and user-friendly tool for maximum likelihood phylogenetic inference. Bioinformatics. 35(21):4453–4455. 10.1093/bioinformatics/btz305

Krogh A, Larsson B, von Heijne G, Sonnhammer EL. 2001. Predicting transmembrane protein topology with a hidden Markov model: application to complete genomes. J Mol Biol. 305(3):567–580. 10.1006/jmbi.2000.4315

Kroll E et al. 2026. The impact of long-read sequencing on fungal genome assemblies: progress and disparity. 2026.05.12.724544 [accessed 2026 Aug 18]. https://www.biorxiv.org/content/10.64898/2026.05.12.724544v1. 10.64898/2026.05.12.724544

Kulik T et al. 2023. Two distinct *Fusarium graminearum* populations colonized European wheat in the past two decades. PLOS ONE. 18(12):e0296302. 10.1371/journal.pone.0296302

Li H. 2018. Minimap2: pairwise alignment for nucleotide sequences. Bioinformatics. 34(18):3094–3100. 10.1093/bioinformatics/bty191

Li H, Durbin R. 2024. Genome assembly in the telomere-to-telomere era. Nat Rev Genet. 25(9):658–670. 10.1038/s41576-024-00718-w

Lovell JT et al. 2022. GENESPACE tracks regions of interest and gene copy number variation across multiple genomes. Elife. 11:e78526. 10.7554/eLife.78526

Lu P et al. 2022. Landscape and regulation of alternative splicing and alternative polyadenylation in a plant pathogenic fungus. New Phytologist. 235(2):674–689. 10.1111/nph.18164

Majidian S et al. 2025. Orthology inference at scale with FastOMA. Nat Methods. 22(2):269–272. 10.1038/s41592-024-02552-8

Manni M et al. 2021. BUSCO Update: Novel and Streamlined Workflows along with Broader and Deeper Phylogenetic Coverage for Scoring of Eukaryotic, Prokaryotic, and Viral Genomes. Mol Biol Evol. 38(10):4647–4654. 10.1093/molbev/msab199

Moolhuijzen PM et al. 2022. A global pangenome for the wheat fungal pathogen *Pyrenophora tritici-repentis* and prediction of effector protein structural homology. Microb Genom. 8(10):mgen000872. 10.1099/mgen.0.000872

Petersen TN, Brunak S, von Heijne G, Nielsen H. 2011. SignalP 4.0: discriminating signal peptides from transmembrane regions. Nat Methods. 8(10):785–786. 10.1038/nmeth.1701

Plissonneau C, Hartmann FE, Croll D. 2018. Pangenome analyses of the wheat pathogen *Zymoseptoria tritici* reveal the structural basis of a highly plastic eukaryotic genome. BMC Biol. 16(1):5. 10.1186/s12915-017-0457-4

Proctor RH, Hohn TM, McCormick SP. 1995. Reduced virulence of *Gibberella zeae* caused by disruption of a trichothecene toxin biosynthetic gene. Mol Plant Microbe Interact. 8(4):593–601. 10.1094/mpmi-8-0593

Richards JK et al. 2018. Reference Quality Genome Assemblies of Three *Parastagonospora nodorum* Isolates Differing in Virulence on Wheat. G3 Genes|Genomes|Genetics. 8(2):393–399. 10.1534/g3.117.300462

Savojardo C, Martelli PL, Fariselli P, Casadio R. 2018. DeepSig: deep learning improves signal peptide detection in proteins. Bioinformatics. 34(10):1690–1696. 10.1093/bioinformatics/btx818

Shen W, Sipos B, Zhao L. 2024. SeqKit2: A Swiss army knife for sequence and alignment processing. iMeta. 3(3):e191. 10.1002/imt2.191

Shumate A, Salzberg SL. 2021. Liftoff: accurate mapping of gene annotations. Bioinformatics. 37(12):1639–1643. 10.1093/bioinformatics/btaa1016

Sieber CMK et al. 2014. The *Fusarium graminearum* Genome Reveals More Secondary Metabolite Gene Clusters and Hints of Horizontal Gene Transfer. PLOS ONE. 9(10):e110311. 10.1371/journal.pone.0110311

Smit AFA, Hubley R, Green P. 2013. RepeatMasker Open-4.0. [accessed 2026 Aug 18]. http://www.repeatmasker.org

Sperschneider J et al. 2016. EffectorP: predicting fungal effector proteins from secretomes using machine learning. New Phytologist. 210(2):743–761. 10.1111/nph.13794

Sperschneider J et al. 2018. Improved prediction of fungal effector proteins from secretomes with EffectorP 2.0. Mol Plant Pathol. 19(9):2094–2110. 10.1111/mpp.12682

Sperschneider J, Dodds PN. 2022. EffectorP 3.0: Prediction of Apoplastic and Cytoplasmic Effectors in Fungi and Oomycetes. Mol Plant Microbe Interact. 35(2):146–156. 10.1094/MPMI-08-21-0201-R

Stanojević D et al. 2026. Telomere-to-telomere assembly using HERRO-corrected Nanopore Simplex reads. Nature. 655(8121):158–165. 10.1038/s41586-026-10563-y

Tettelin H et al. 2005. Genome analysis of multiple pathogenic isolates of *Streptococcus agalactiae*: Implications for the microbial “pan-genome.” Proceedings of the National Academy of Sciences. 102(39):13950–13955. 10.1073/pnas.0506758102

Teufel F et al. 2022. SignalP 6.0 predicts all five types of signal peptides using protein language models. Nat Biotechnol. 40(7):1023–1025. 10.1038/s41587-021-01156-3

Thumuluri V et al. 2022. DeepLoc 2.0: multi-label subcellular localization prediction using protein language models. Nucleic Acids Res. 50(W1):W228–W234. 10.1093/nar/gkac278

Uliano-Silva M et al. 2023. MitoHiFi: a python pipeline for mitochondrial genome assembly from PacBio high fidelity reads. BMC Bioinformatics. 24(1):288. 10.1186/s12859-023-05385-y

Urban M et al. 2016. First Draft Genome Sequence of a UK Strain (UK99) of *Fusarium culmorum*. Genome Announc. 4(5):e00771–16. 10.1128/genomeA.00771-16

Urban M et al. 2022. PHI-base in 2022: a multi-species phenotype database for Pathogen–Host Interactions. Nucleic Acids Research. 50(D1):D837–D847. 10.1093/nar/gkab1037

Urban M et al. 2025. PHI-base - the multi-species pathogen-host interaction database in 2025. Nucleic Acids Res. 53(D1):D826–D838. 10.1093/nar/gkae1084

Venturini L et al. 2018. Leveraging multiple transcriptome assembly methods for improved gene structure annotation. Gigascience. 7(8):giy093. 10.1093/gigascience/giy093

Walkowiak S, Rowland O, Rodrigue N, Subramaniam R. 2016. Whole genome sequencing and comparative genomics of closely related Fusarium Head Blight fungi: *Fusarium graminearum, F. meridionale* and *F. asiaticum*. BMC Genomics. 17(1):1014. 10.1186/s12864-016-3371-1

Wang Q et al. 2017. Characterization of the Two-Speed Subgenomes of *Fusarium graminearum* Reveals the Fast-Speed Subgenome Specialized for Adaption and Infection. Front Plant Sci. 8. 10.3389/fpls.2017.00140

Wang Y et al. 2012. MCScanX: a toolkit for detection and evolutionary analysis of gene synteny and collinearity. Nucleic Acids Res. 40(7):e49. 10.1093/nar/gkr1293

Wood AK et al. 2020. Genome Sequence of *Fusarium graminearum* Strain CML3066, Isolated from a Wheat Spike in Southern Brazil. Microbiol Resour Announc. 9(19):e00157–20. 10.1128/MRA.00157-20

Wyatt NA, Richards JK, Brueggeman RS, Friesen TL. 2020. A Comparative Genomic Analysis of the Barley Pathogen *Pyrenophora teres f. teres* Identifies Subtelomeric Regions as Drivers of Virulence. MPMI. 33(2):173–188. 10.1094/MPMI-05-19-0128-R

Zheng J et al. 2023. dbCAN3: automated carbohydrate-active enzyme and substrate annotation. Nucleic Acids Res. 51(W1):W115–W121. 10.1093/nar/gkad328

